# Legacy 4(1*H*)-quinolone scaffolds activity against acute and chronic *Toxoplasma gondii* infection

**DOI:** 10.64898/2026.03.10.710892

**Authors:** Melissa A. Sleda, Khaly Diagne, Victoria M. Clifton, Baihetiya Baierna, Roman Manetsch, Silvia NJ Moreno

## Abstract

*Toxoplasma gondii* is a protozoan parasite capable of infecting most warm-blooded animals, including humans, and can cause severe disease in immunocompromised individuals and the developing fetus. Current treatments for toxoplasmosis are effective only against the acute stage of infection and have limited or no activity against the latent bradyzoite stage found within tissue cysts. The mitochondrion of *T. gondii* is a validated drug target, and the clinically used drug atovaquone acts by inhibiting the mitochondrial electron transport chain (ETC) at the coenzyme Q:cytochrome c oxidoreductase (*bc_1_* complex). In this study, we evaluate two legacy 4(1*H*)-quinolones: ICI 56,780 and WR 243246, previously shown to inhibit the *Plasmodium falciparum bc_1_* complex, for their efficacy against *T. gondii*. Both compounds inhibit tachyzoite growth with low-nanomolar EC₅₀ values and disrupt parasite mitochondrial function by blocking cytochrome *c* reduction and collapsing the mitochondrial membrane potential. Notably, ICI 56,780 protects mice from lethal infection with type I RH tachyzoites. Importantly, ICI 56,780 also exhibits potent activity against chronic-stage parasites, reducing cyst size and bradyzoite viability *in vitro* and showing low-nanomolar EC₅₀ values against *in vivo*-derived bradyzoites. In mice chronically infected with *T. gondii*, treatment with ICI 56,780 significantly decreases brain cyst burden. Although these 4(1*H*)-quinolones display some pharmacokinetic limitations, our findings highlight their potential as promising chemotypes active against both acute and chronic stages of *T. gondii* and provide a framework for future medicinal chemistry efforts to improve drug-like properties while preserving or enhancing anti-bradyzoite activity.

## Introduction

*Toxoplasma gondii* is an obligate intracellular parasite that infects approximately one third of the global human population^1^. Upon infection, *T. gondii* replicates as tachyzoites, the fast-replicating form that can destroy host tissues and disseminate throughout the body. In immunocompetent individuals, the immune response controls the acute infection, and tachyzoites subsequently differentiate into bradyzoites, a slow-growing form that resides within tissue cysts. These cysts can persist in the host for life, mainly in the brain and muscle tissues^2^. However, if an individual becomes immunosuppressed, bradyzoites can reactivate reverting to tachyzoites and cause serious disease. Current treatments for toxoplasmosis, including pyrimethamine and sulfadiazine, are effective against tachyzoites^3^ but have minimal activity against the bradyzoite stage within tissue cysts^4, 5^. The last clinically approved drug, atovaquone (ATQ), targets the mitochondrial *bc_1_* complex^6, 7^ and shows some effect against bradyzoites, but is unable to completely eliminate tissue cysts of the chronic infection^8^.

Mitochondrial metabolism has emerged as an important vulnerability of *T. gondii* for both acute and chronic stages of infection. Although bradyzoites were long thought to exhibit reduced metabolic activity compared with tachyzoites, raising questions about the extent of mitochondrial metabolism in this stage^9, 10^, several studies have shown that mitochondrial pathways remain functionally relevant during the chronic stage^11^. Consistent with this, the cytochrome *bc_1_* inhibitor ATQ, reduces tissue cyst numbers and mitigates neuropathology in chronically infected mice^8, 12^. More recent research further validates the mitochondrion as a viable target during bradyzoite development^13, 14^.

Our laboratory identified a lipophilic bisphosphonate that inhibits ubiquinone (UQ) synthesis, an essential component of the ETC, and demonstrated that it is highly effective at protecting mice against a lethal infection with *T. gondii*^15^, while significantly reducing tissue cyst burden^16^. Other inhibitors of the *bc_1_* complex, like endochin-like quinolones (ELQs), similarly decrease tissue cyst burden in chronically infected mice^17–19^. Together, these findings underscore the sensitivity of bradyzoites to mitochondrial disruption and highlight the ETC as a promising target for therapeutic intervention.

Quinolone compounds have been shown to inhibit DNA replication by targeting DNA gyrase and topoisomerase IV^20^. In addition, quinolone derivatives have also been reported to induce apoptosis in cancer cells^21, 22^, modulate bacterial redox homeostasis^23^, and chelate metal ions, which can alter enzymatic functions and metabolic processes^24^. The 4(1*H*)-quinolone scaffold has re-emerged as a promising chemotype for antiparasitic drug development, with historical roots in the antimalarial compound ICI 56,780 (Fig. 1A). Initially discovered in the mid-20^th^ century, ICI 56,780 demonstrated potent activity against *Plasmodium* spp. but was ultimately discontinued due to poor aqueous solubility and rapid emergence of resistance *in vivo*^25^. Despite these limitations, its ability to inhibit the parasite mitochondrial cytochrome *bc₁* complex provided a valuable pharmacological foundation. Extensive structure-activity (SAR) and structure-property (SSR) relationship studies by Manetsch and colleagues refined this scaffold, yielding analogs with nanomolar potency against multidrug-resistant *Plasmodium falciparum*, *in vivo* efficacy across parasite life stages, and *bc₁*complex inhibition as the confirmed mechanism of action^26,27–29,30,25,31^. To improve metabolic and physicochemical properties, 7-(2-phenoxyethoxy)- and 7-piperazine-substituted 4(1*H*)-quinolones were synthesized and shown to possess nanomolar potency against multidrug-resistant *P. falciparum*, alongside robust *in vivo* efficacy^25, 31^. These optimized analogs retained activity against both blood- and liver-stage parasites and displayed oral bioavailability and curative potential in murine malaria models. Mechanistic studies confirmed that these compounds act via inhibition of the cytochrome *bc₁* complex, consistent with their structural lineage. Given the conserved mitochondrial ETC machinery across apicomplexan parasites, this class of compounds warrants further investigation against related pathogens such as *T. gondii*.

**Figure 1.**
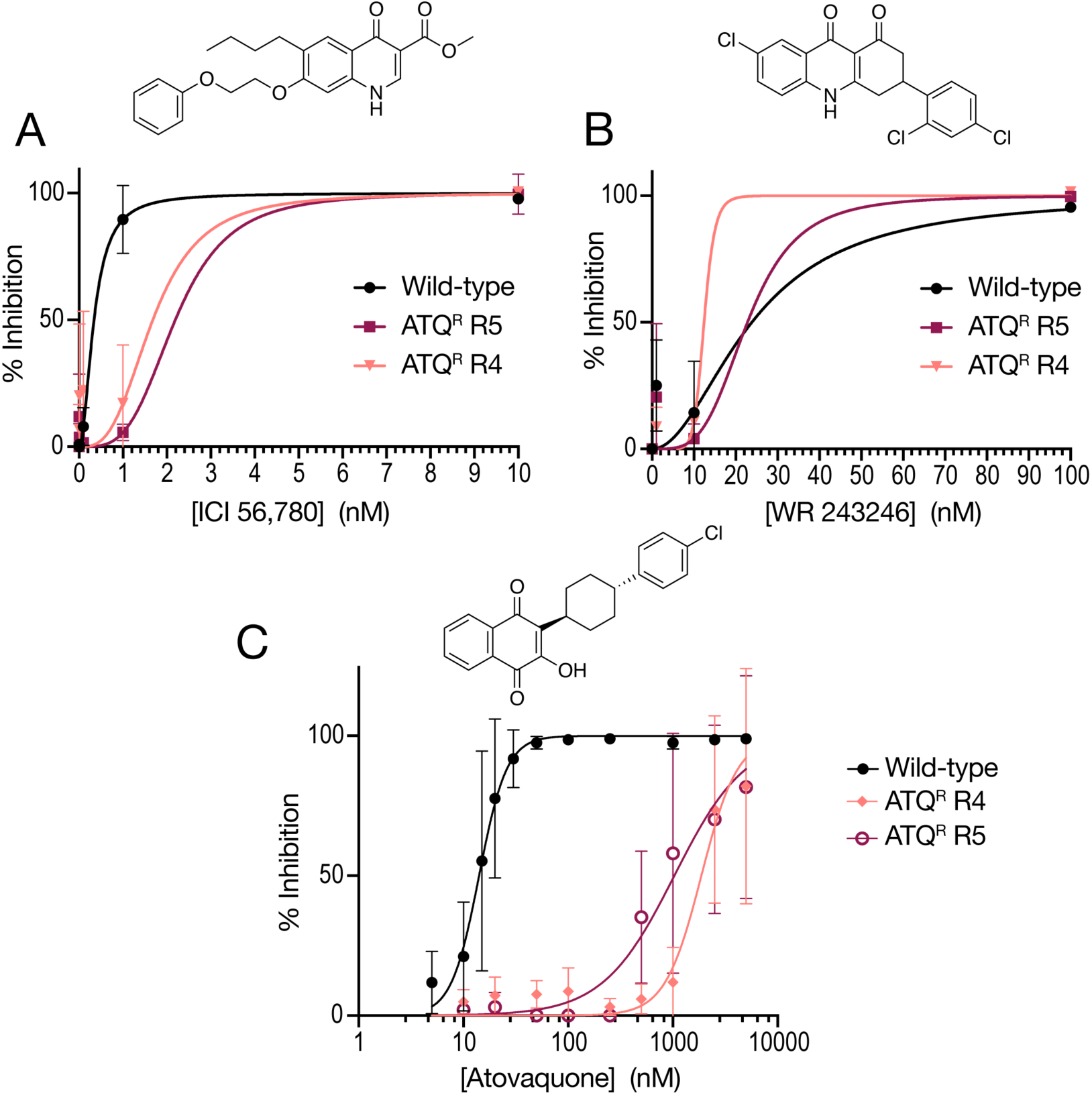
Inhibition of tachyzoite growth by ICI 56,780 and WR 243246. A. EC_50_ curves for ICI 56,780 in RH-RFP (type I) wild-type tachyzoites, ME49 (type II) ATQ^R^ strains R4 and R5. Growth was measured by evaluating red fluorescence of RH-RFP parasites or by quantifying crystal violet absorbance for the R4 and R5 strains. B. EC_50_ curves for WR 243246 in RH-RFP and ATQ^R^ strains R4 and R5. C. EC_50_ curves for ATQ in RH-RFP and ATQ^R^ strains R4 and R5. Each curve represents the average from 3 biological replicates. A non-linear regression was done to calculate EC_50_ using prism.

The dihydroacridinedione scaffold has also contributed to antimalarial drug discovery, with WR 243246 serving as an early lead compound. Further optimization of 1,2,3,4-tetrahydroacridin-9(10H)-ones (THAs) produced derivatives with enhanced potency and improved pharmacokinetic profiles, including efficacy against chloroquine-resistant *P. falciparum* strains^27^. These findings support the exploration of THA analogs against other apicomplexan pathogens.

The mitochondrion of *T. gondii* has emerged as a promising target for therapeutic intervention, given its essential role in parasite survival and its divergence from mammalian host mitochondria^32, 33^. Several studies have demonstrated that inhibitors targeting the mitochondrial ETC, such as ATQ and ELQs, reduce tissue cyst burden in murine models, underscoring the ETC’s critical function in bradyzoite viability^19^. Additional mitochondrial enzymes with no mammalian homologs, such as malate:quinone oxidoreductase (MQO)^34^, provide opportunities for selective targeting. The identification of mitochondrial proteins such as TgPRELID, which has been linked to resistance to mitochondrial inhibitors, further highlights the importance of mitochondrial pathways in parasite physiology and drug response^35^. Collectively, these findings support continued exploration of mitochondrial pathways as promising targets for developing effective treatments against chronic toxoplasmosis.

In this study, we explore the efficacy of two 4(1*H*)-quinolones, ICI 56,780 and WR 243246, against *T. gondii*. Both ICI 56,780 and WR 243246 differ from the ELQ quinolone series in the position of the main substituent in the quinolone core. We investigate their impact on both tachyzoite and bradyzoite stages, assess their mechanism of action, and evaluate their effectiveness against strains resistant to ATQ. These findings contribute to the development of more effective therapies targeting chronic *T. gondii* infection.

## Results

### Inhibition of T. gondii Tachyzoite Proliferation and Mitochondrial Function

We tested the activity of the legacy 4(1*H*)-quinolones ICI 56,780 and WR 243246 against *in vitro* proliferation of *T. gondii* tachyzoites of the type I RH strain and two Type II ATQ resistant (ATQ^R^) strains (R4 and R5)^7^. We found that ICI 56,780 inhibits *T. gondii* RH with an EC_50_ of 0.34 nM and it retained efficacy against both ATQ^R^ strains with EC_50_ values of 1.68 nM and 2.19 nM against R4 and R5, respectively (Fig. 1A, Table 1). WR 243246 is also effective against RH with an EC_50_ of 24.2 nM and is equally effective against R4 and R5 with EC_50_ values of 22.7 nM and 12.5 nM, respectively (Fig. 1B, Table 1). For comparison, ATQ displayed an EC_50_ of 14.1 nM against wild-type RH, but resistance in the R4 and R5 strains resulted in a marked increased EC_50_s to 1.87 μM and 1.03 μM, respectively (Fig. 1C).

**Table 1.**
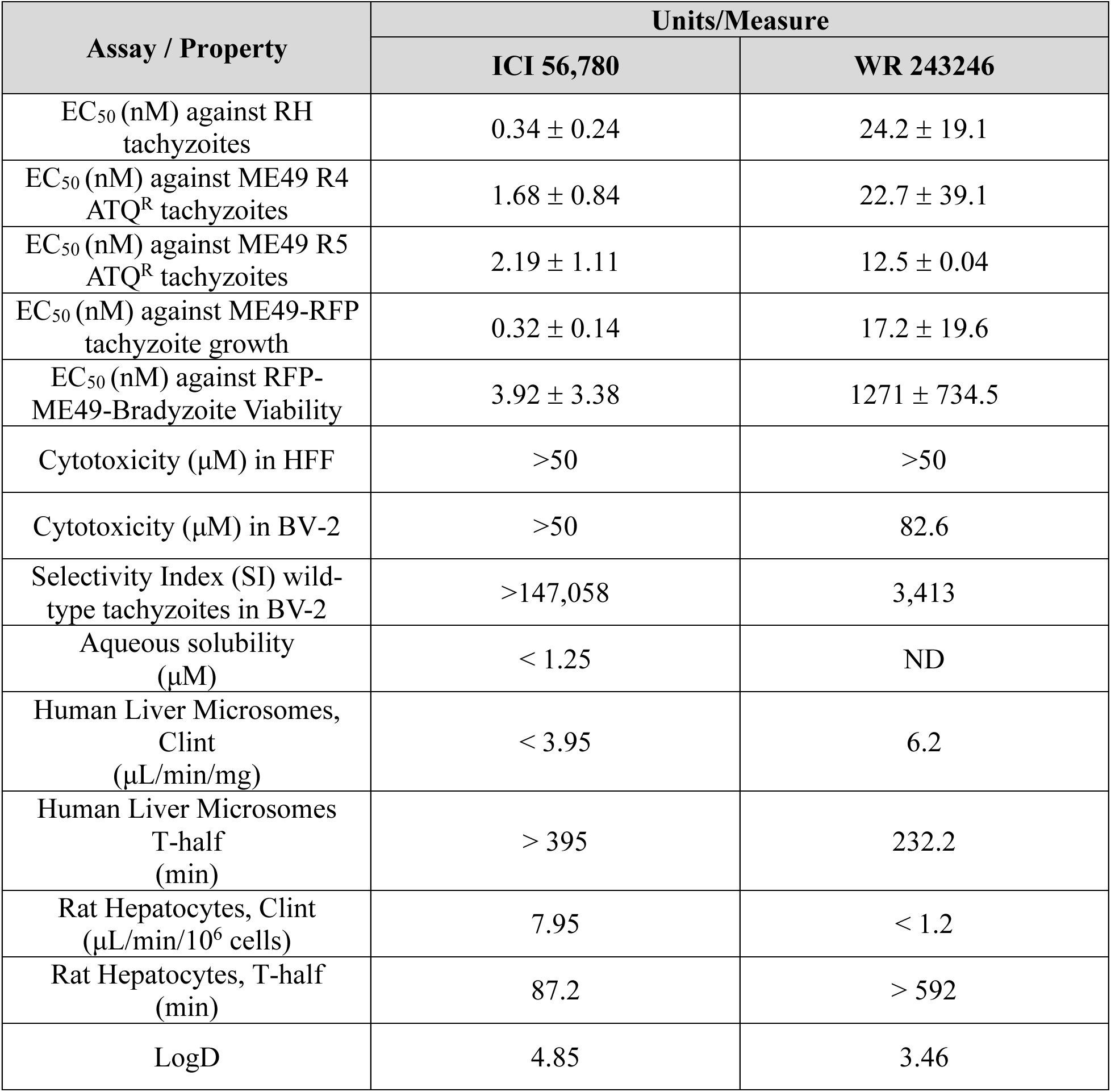
Inhibitory concentrations EC_50_s against tachyzoites and bradyzoites, cytotoxicity in host cells (fibroblasts and glial cells), and physiochemical properties of ICI 56,780 and WR 243246.

To investigate the effects of ICI 56,780 and WR 243246 on mitochondrial functions, we performed a succinate cytochrome c reductase assay, which assesses the activity of both Complex II and III, using succinate as the substrate (Fig. 2A). Both compounds inhibited cytochrome c reduction similarly to ATQ, confirming ETC inhibition (Fig. 2B).

**Figure 2.**
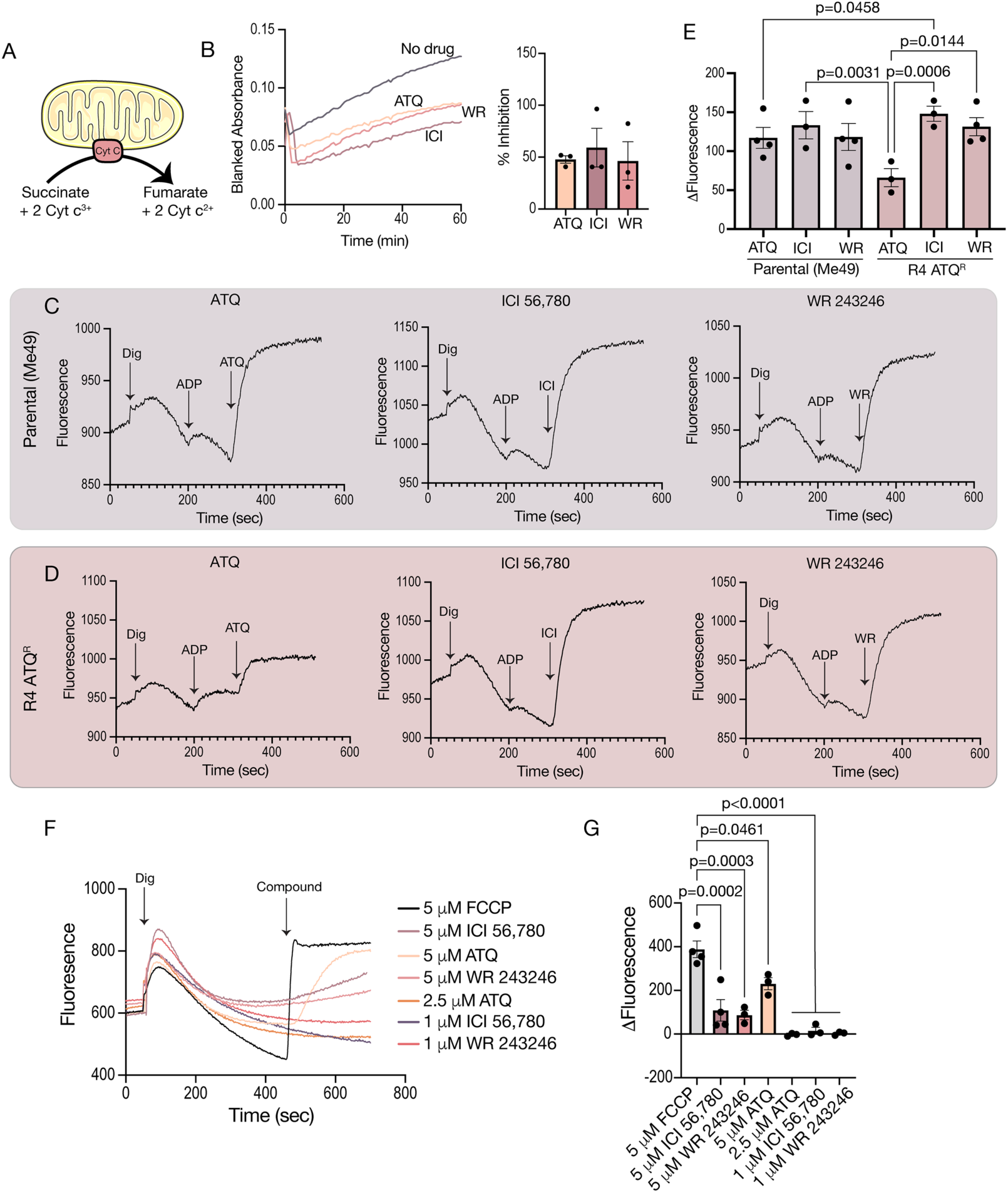
Effects of ICI 56,780 and WR 243246 on Mitochondrial Electron Transport and Membrane Potential. A. Schematics of the cytochrome c reduction assay. B. Representative traces for the cytochrome c reduction activity of *T. gondii* mitochondrial fractions without any additions or in the presence of the indicated inhibitors at 100 nM. The bar graph shows the averages from 3 independent biological replicates. C. Representative traces of mitochondrial membrane potential measured with Safranin O (2.5 μM) in the parental cell line (ME49) during real-time addition of the indicated compounds (Dig: Digitonin, 10 μM; ADP, 10 μM). D. Representative traces of mitochondrial membrane potential in the R4 ATQ^R^ cell line during real-time addition of each drug. E. Quantification of the average change in fluorescence after each addition (2.5 μM ATQ; 1 μM ICI 56,780 and WR 243246), from at least three independent biological replicates. F. Representative tracings of hTert fibroblast mitochondrial membrane potential measurements. Digitonin (50 μM) was added at 50 sec and compounds (at indicated concentrations) were added at 450 sec. FCCP at 5 μM was used as the control for maximum depolarization. G. Quantification of the average change in fluorescence after drug addition, from 3 independent biological replicates. Statistical analysis in E and G were performed using two-way ANOVA.

Additionally, we analyzed the mitochondrial membrane potential (ΔΨ_m_) using safranin O in digitonin-permeabilized tachyzoites in the presence of succinate as substrate^36^. Following digitonin permeabilization, Safranin O accumulates within polarized mitochondria, leading to a characteristic decrease in fluorescence that reflects the membrane potential. Upon addition of ADP, the ATP synthase is activated, which utilizes the proton gradient and induces partial mitochondrial depolarization. This depolarization leads to the redistribution of safranin O resulting in a corresponding change in fluorescence. To evaluate the effect of the test compounds on mitochondrial membrane potential, we added them at 1 μM and monitored the resulting changes in Safranin O fluorescence. Both ICI 56,780 and WR 243246 depolarized the mitochondrial membrane potential similarly to the ATQ used as control (Fig. 2C-E). In the ATQ^R^ cell line, the fluorescence changes normally induced by ATQ was markedly reduced. However, both ICI 56,780 and WR 243246 remained equally effective in the ATQ^R^ strain compared to the parental ME49 strain (Fig. 2E). This suggests that, although these compounds inhibit the ETC like ATQ, they likely target a different site within the cytochrome *bc₁* complex.

Additionally, we tested the effects of these compounds on the membrane potential of mammalian host cells (hTERT fibroblasts) at the same concentrations that inhibit parasite growth. No significant change in membrane potential was observed under conditions similar to those used for the parasites (Fig. 2F-G). However, when the concentration of each compound was increased to 5 µM, a significant reduction in host mitochondrial membrane potential was detected (Fig. 2F-G). FCCP was used as a positive control and defined as 100% inhibition of membrane potential^37^. These results indicate that the compounds do not cause detectable toxicity to host mitochondria at concentrations that inhibit parasite growth.

### Physicochemical properties and compound stability

Considering that we aimed to evaluate these compounds in both mouse models of toxoplasmosis, acute and chronic, we next profiled key physicochemical and metabolic properties of ICI 56,780 and WR 243246 to identify potential liabilities that might limit efficacy studies in animals. Both compounds are lipophilic and poorly soluble in aqueous media, with logD values of 4.85 for ICI 56,780 and 3.5 for WR 243246. The aqueous solubility of ICI 56,780 was < 2 µM; while the solubility of WR 243246 was too low to be reliably determined in the solubility assay used. Both compounds are also predicted to exhibit high plasma protein binding, which further limits their suitability for detailed pharmacokinetic profiling in mice at this stage. Consequently, future structure-activity relationship optimization will place strong emphasis on improving the physicochemical properties of these chemotypes to enable oral delivery and to achieve plasma concentrations sufficient to produce robust *in vivo* efficacy. Metabolic stability was determined using rat hepatocytes and human liver microsomes, with rat serving as a rodent comparator commonly used in preclinical studies and human microsomes providing translational insight into potential human metabolic stability. ICI 56,780 exhibited good metabolic stability in both human microsomes and in rat hepatocytes with a very low intrinsic clearance (Clint values ≤ 8 μL/min/10^6^ cells and half-life values ≥ 90 minutes). WR 243246, was similarly stable in human microsomes (Clint = 6.2 μL/min/mg, t_1/2_ >592 min), but was significantly less stable in rat hepatocytes, with a very short half-life of less than 2 min (Table 1) indicating species-specific metabolic instability. Importantly, these profiling studies indicate that while the basic physicochemical properties of both ICI 56,780 and WR 243246 are not optimal, they are sufficient to support *in vivo* efficacy studies.

We also evaluated host cytotoxicity in HFF fibroblasts and BV-2 microglial cells. No evidence of toxicity was observed for either compound in HFF fibroblasts (Table 1 and supplementary Table 1). In BV-2 microglia, however, cytotoxicity was observed for ATQ (CC_50_ = 33.24 μM) and WR 243246 (CC_50_ = 82.6 μM) (Table 1 and supplementary Table 2). No evidence of toxicity was detected for ICI 56,780 preventing determination of a CC_50_ (Supplementary Table 2).

### Activity of ICI 56,780 and WR 243246 in the acute infection model

We next evaluated *in vivo* activity using an acute infection model in Swiss Webster mice. Female Swiss Webster mice were infected intraperitoneally (i.p.) with 100 RH-RFP *T. gondii* tachyzoites, and treatment with the test compounds was initiated 6 hours post-infection at the indicated doses. Treatment continued for 10 consecutive days, during which body weight and survival were monitored (Fig. 3A). ICI 56,780 protected mice from lethal infection at doses as low as 1 mg/kg/day and provided complete protection at 10 mg/kg/day (Fig. 3B). For comparison, atovaquone (ATQ) has been reported to result in 100% mortality at 4.6 mg/kg requiring doses of 75 mg/kg to achieve full protection in this model^38^. Weight monitoring revealed that treated mice experienced transient weight loss around days 8-12, coinciding with visible signs of illness. However, the mice recovered and regained weight (Supplementary Figure 1, A-C). To verify that the surviving mice had been infected and had mounted an immune response, we collected serum and tested for *T. gondii*-specific antibodies at 28 days post-infection (dpi), prior to challenge (Supplementary Figure 1, D-F). At 30 days post-infection, surviving mice were challenged with 10,000 RH-RFP tachyzoites and body weight and survival was monitored for three weeks before the animals were euthanized. We also tested WR 243246 in the acute lethal infection model. However, only a single mouse treated with 4 mg/kg/day survived the infection, while all other treated animals succumbed to infection (Fig. 3C; Supplementary Figure 1G,I). Increasing the doses to 5, 10 and 20 mg/kg/day did not improve the survival outcome (Fig. 3D, Supplementary Figure 1H). This lack of efficacy at higher doses may reflect limited bioavailability of WR 243246, an issue that we plan to address through development of improved analogs.

**Figure 3.**
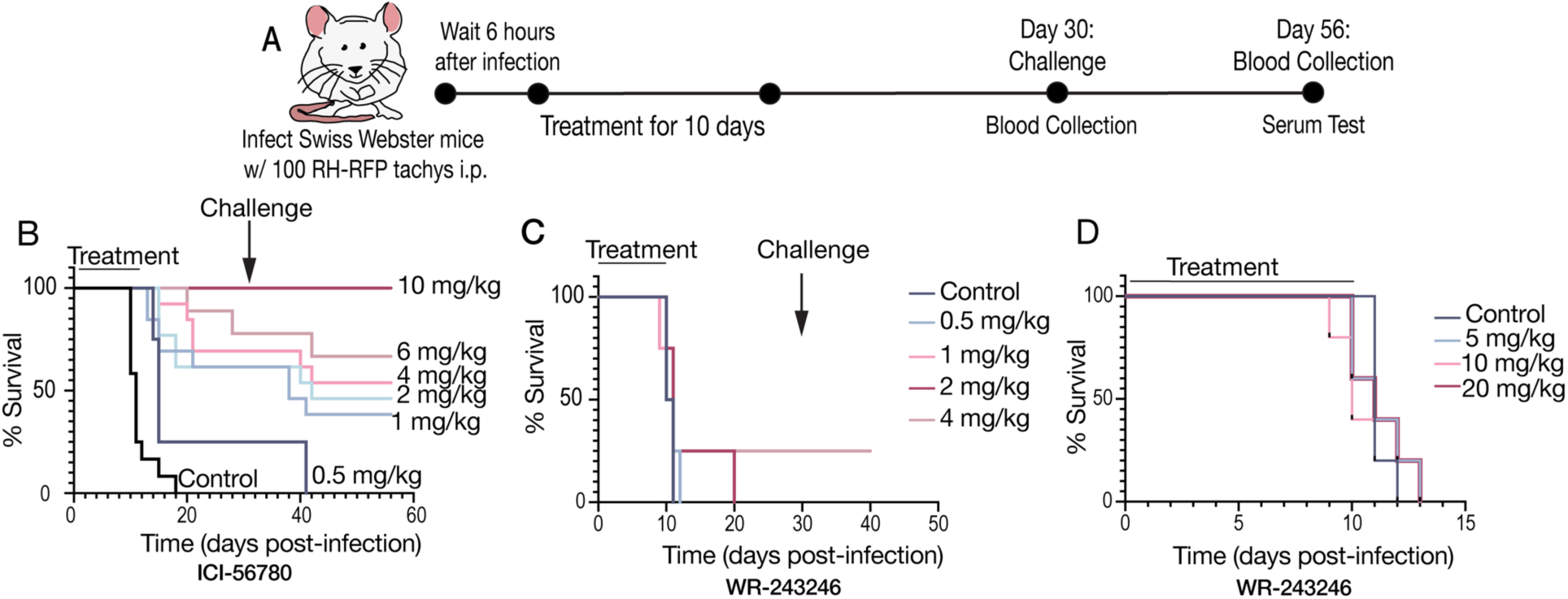
Acute *T. gondii* infection model and treatment in mice. A. Schematic of the acute infection model. Swiss webster mice were infected with 100 RH-RFP tachyzoites i.p.. Treatment was initiated 6 hours post-infection and administered daily by i.p. injection at the doses indicated in the graphs. At 30 dpi blood was collected for serum testing and mice were challenged with 10,000 RH-RFP tachyzoites. Three weeks after the challenge mice were euthanized and the final blood was collected for serum testing. B. Survival curve for 3 biological replicates of mice treated with ICI 56,780 with 0.5, 1, 2, 4, 6, and 10 mg/kg/day. C. Survival curve of one biological replicate of mice treated with WR 243246 at 0.5, 1, 2, and 4 mg/kg. D. Survival curve for one biological replicate of mice treated with WR 243246 at 5, 10, and 20 mg/kg.

### Activity of ICI 56,780 and WR 243246 against in vitro-derived bradyzoites

To determine whether the compounds also affect the chronic stage, we evaluated their activity against bradyzoites. We first tested them against *in vitro* derived bradyzoites using our previously established protocols to differentiate ME49 using compound 1 (4-[2-(4-fluorophenyl)-5-(1-methylpiperidine-4-yl)-1H-pyrrol-3-yl]pyridine) and low CO_2_ growth conditions^39^. After differentiation of tachyzoites into bradyzoites, the cultures were treated with 30 nM of each compound for 4 days. We used fluorescence lectin staining with FITC-conjugated Dolichos biflorus agglutinin (DBA) to visualize tissue cysts, as DBA labels the cyst wall^40^ together with an immunofluorescence assay for BAG1 (bradyzoite antigen 1). Additionally, we stained the cysts with anti-SAG1 (tachyzoite surface antigen 1), whose expression indicates that the *in vitro* derived cysts are immature (Fig. 4A-B). We found that both ICI 56,780 and WR 243246 reduced cyst size (Fig. 4B). Notably, WR 243246-treated cultures showed internal spaces inside the vacuoles not occupied by parasites indicating disruption of cyst morphology (Fig. 4B). We next tested bradyzoite viability by mechanically disrupting the *in vitro*-derived cysts, followed by pepsin treatment to release bradyzoites and eliminate any remaining tachyzoites. The recovered bradyzoites were then used in plaque assays to evaluate their viability (Fig. 4C). Under these conditions, surviving bradyzoites convert to tachyzoites and initiate lytic cycles that disrupt the host monolayer, forming characteristic plaques. The number of plaques formed reflects the number of initial viable bradyzoites present in the sample.

**Figure 4.**
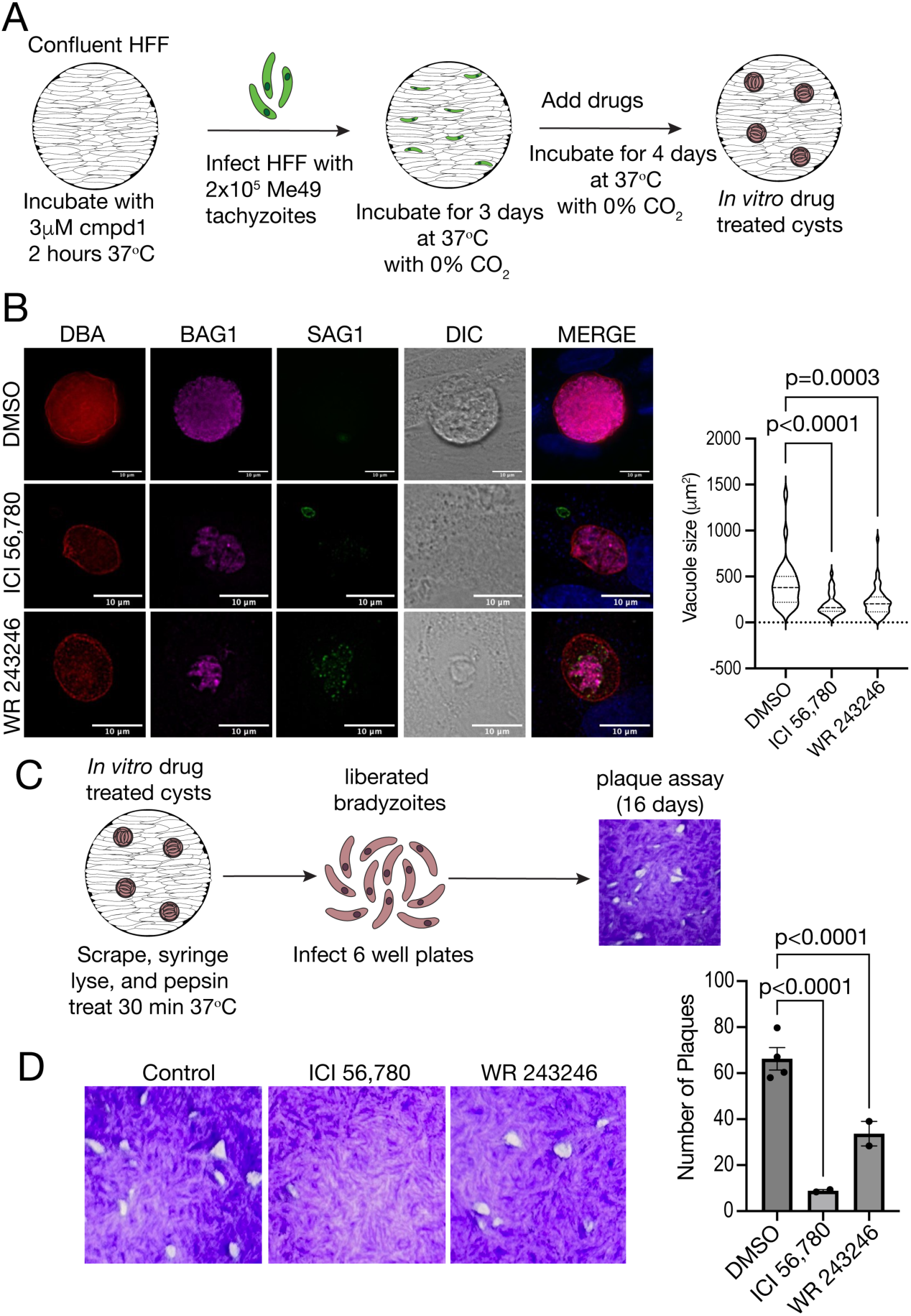
Activity of ICI 56,780 and WR 243246 against *in vitro*-derived bradyzoites. A. Schematic showing the protocol for *in vitro* differentiation of ME49 tachyzoites into bradyzoites. HFF monolayers were incubated with compound 1, 2 hours prior to infection with 5,000 ME49 tachyzoites. The cells were incubated for 3 days at ambient CO_2_. Compounds were then added at 30 nM and incubated for an additional 4 days at ambient CO_2_ before fixation and staining with markers for the cyst wall (DBA), bradyzoites (BAG1) and tachyzoites (SAG1). B. Representative images for each condition, scale bar is 10 μm. The size of the cysts was measured using ImageJ and averaged from 3 biological replicates. C. Schematic showing the protocol used for the *in vitro* bradyzoite viability assay. First, tachyzoites were differentiated as described in A, then the drug treated cysts were syringe lysed and treated with pepsin to release bradyzoites. The bradyzoites are then plated onto confluent HFF monolayers at 10,000 bradyzoites per well and allowed to grow under normal growth conditions with drug-free DMEM for 16 days. D. Representative images of plaques for each condition. The bar graph shows the average number of plaques formed from three independent biological replicates. Statistical significance was assessed using two-way ANOVA, where indicated (B and D).

There was a significant decrease in the number of plaques formed by both ICI 56,780 and WR 243246 treated bradyzoites indicating that both compounds are effective against the viability of *in vitro* derived bradyzoites (Fig. 4D).

### Activity of ICI 56,780 and WR 243246 against in vivo-derived bradyzoites

We also tested the compounds against *in vivo* derived bradyzoites given the known differences between *in vitro* and *in vivo* derived tissue cysts^41^. Using a previously established *ex vivo* protocol adapted from Radke et al.^42^, we assessed the ability of the compounds on the viability of bradyzoites isolated from chronically infected mice. Tissue cysts were isolated from the brains of chronically infected mice by homogenization followed by acid/pepsin treatment to release bradyzoites. The recovered bradyzoites were then used in two assays: continuous drug treatment to assess inhibition of ME49-RFP tachyzoite growth and short (4-h) exposure to evaluate effects on bradyzoite viability (Fig. 5A).

**Figure 5.**
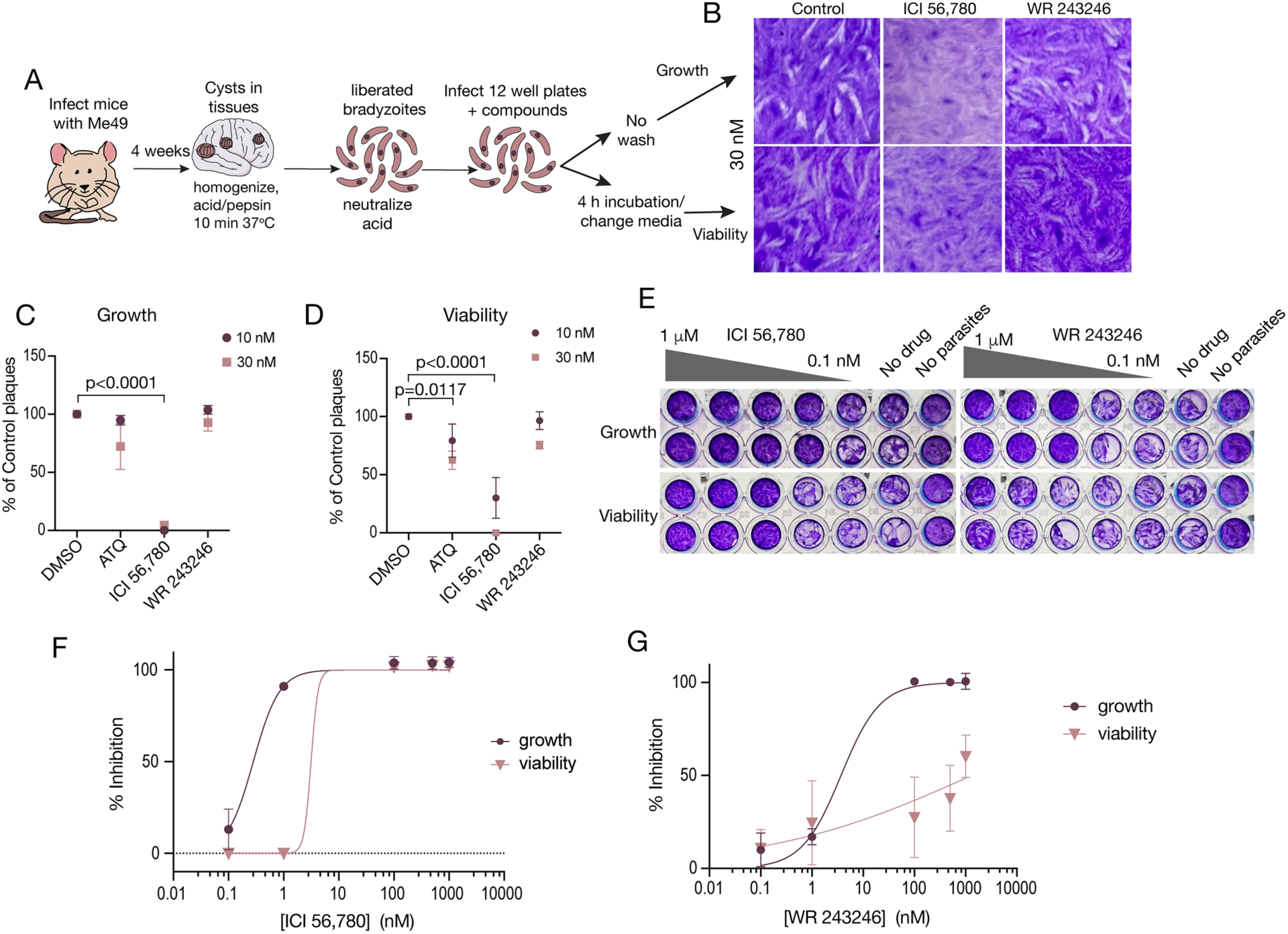
ICI 56,780 and WR 243246 inhibition of *in vivo*-derived bradyzoites. A. Schematic of the protocol used for the *ex vivo* assay. Mice are infected with ME49 tachyzoites, and four weeks post-infection brains are collected, homogenized and cysts enumerated. Cysts are treated with acid/pepsin to liberate bradyzoites, which are used for two different assays: a growth assay in which parasites are continuously exposed to the compounds for 14 days, and a viability assay, in which bradyzoites are treated with compounds for 4 h, followed by washing and incubation in drug-free DMEM. B. Representative images of plaques formed following the viability and growth assays. C. Average number of plaques formed in the growth assay, normalized to the DMSO treated bradyzoites controls, calculated from 3 biological replicates. D. Average number of plaques formed in the viability assay, normalized to the DMSO treated bradyzoites controls, calculated from 3 biological replicates. E. Representative plaques formed in the 96-well plate *ex vivo* assays for both growth and viability, showing dose-response for ICI 56,780 and WR 243246. F. Dose response curves for ICI 56,780 in growth and viability assays from 2 independent biological replicates. G. Dose response curves for WR 243246 in growth and viability assays from 2 independent biological replicates. EC_50_ values were determined by non-linear regression analysis using GraphPad Prism. Statistical significance was assessed using two-way ANOVA, where indicated.

We found that only ICI 56,780 was effective at 30 nM in inhibiting both growth and viability of *in vivo*-derived bradyzoites (Fig. 5B-D). Inhibition of bradyzoite viability provides a more direct and stage-specific measure of chronic-stage activity than inhibition of overall growth, which may primarily reflect effects on reactivated tachyzoites. We next developed a 96-well plate assay to quantify the inhibition and determine EC_50_ values. ICI 56,780 showed exceptional potency, inhibiting bradyzoite viability with an EC_50_ of 3.92 nM, only modestly higher than the EC_50_ measured for growth (0.32 nM) and still well within the low nanomolar range, indicating strong efficacy against both stages (Fig. 5E-F; Table 1). The growth EC_50_ against ME49 tachyzoites (0.32 nM) closely matches the EC_50_ against RH tachyzoites (0.34 nM), indicating that ICI 56,780 is highly effective against tachyzoites of both strains. This high potency, together with its low-nanomolar activity against bradyzoites, highlights its broad efficacy across parasite stages.

WR 243246 also inhibited ME49 tachyzoite growth effectively (EC_50_ = 3.37 nM), but its activity against bradyzoite viability was markedly reduced (EC_50_ = 1271 nM), although still measurable (Fig. 5E, G; Table 1). Thus, while WR 243246 remains active, its reduced potency toward bradyzoites contrasts with the broad, stage-spanning efficacy observed with ICI 56,780.

Altogether, these findings demonstrate that ICI 56,780 is highly effective against both *in vitro* and *in vivo* derived bradyzoites. In contrast, WR 243246 is less effective against *in vivo* derived bradyzoites despite its *in vitro* potency. These results underscore the importance of testing *ex vivo* bradyzoites, to validate compound efficacy before proceeding to *in vivo* studies of chronic infection.

### ICI 56,780 Reduces Brain Cyst Burden in Chronically Infected Mice

Since ICI 56,780 showed strong activity against bradyzoite viability, we next evaluated its efficacy in an *in vivo* model of chronic *T. gondii* infection using CBA/j mice, following a previously established protocol^16^, adapted from Doggett et al.^17^. We infected male CBA/j mice with 1,000 ME49-RFP tachyzoites i.p. and allowed the infection to progress to the chronic stage over 4 weeks. After confirming the presence of tissue cysts in two randomly selected mice, we started daily treatment for 16 days. Following a 2-week recovery period, brains were collected to enumerate tissue cysts (Fig. 6A). Mice were treated with ICI 56,780 at 1 and 5 mg/kg/day, and with ATQ at 5 mg/kg/day included as a positive control. Representative images of tissue cysts stained with DBA and BAG1 are shown in Fig. 6B.

**Figure 6.**
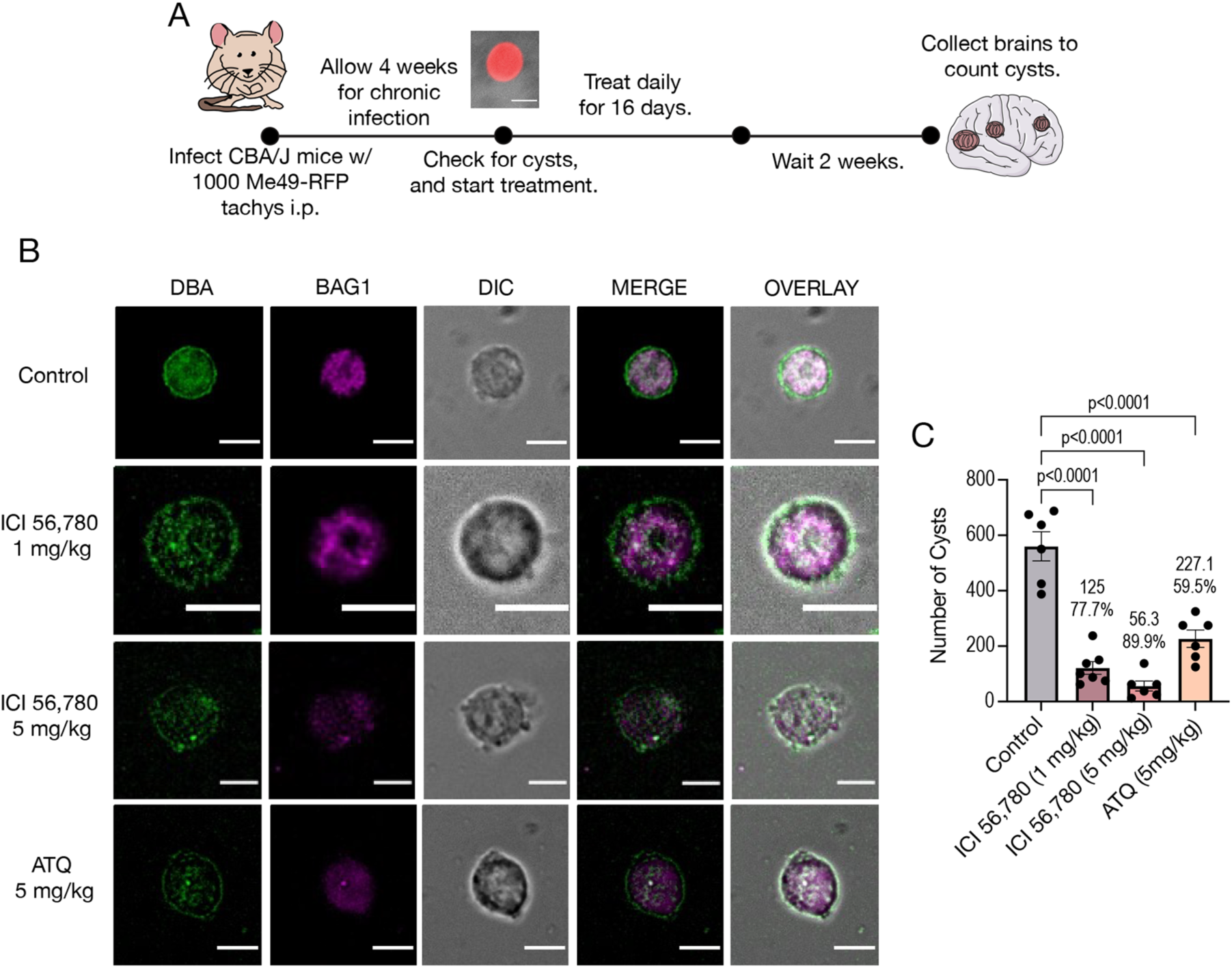
ICI 56,780 reduces brain cyst burden in a mouse model of chronic toxoplasmosis. A. Schematic showing the protocol followed for the *in vivo* chronic infection. CBA/j mice were infected with 1,000 Me49-RFP tachyzoites i.p. Four weeks post-infection, 2 mice were euthanized to collect their brains to confirm the presence of tissue cysts and the remaining mice were treated daily by i.p injection for 16 days with ICI 56,780 dissolved in Kolliphor HS-15 at doses of 1 mg/kg/day and 5 mg/kg/day. ATQ was included as positive control at a dose of 5 mg/kg/day. B. Representative images of cysts visualized with DBA staining and antibodies against BAG1 (bradyzoite surface antigen 1) protein to label cysts and bradyzoites, respectively, from brains of chronically infected mice treated with the indicated drugs. Scale bar=5 μm. C. Quantification of the number of cysts obtained from brains of chronically infected and treated mice. Data represent the mean from two independent biological replicates. Each dot represents a mouse, and each group had 6 mice except the ICI 56,780 1 mg/kg group which had 7 mice total. Statistical significance was assessed using two-way ANOVA, where indicated.

ICI 56,780 significantly reduced brain cyst burden by 77.7% at 1 mg/kg and 89.9% at 5 mg/kg, substantially exceeding the reduction achieved with the ATQ control (59.5%), which is consistent with the known limited efficacy of ATQ against *in vivo* cysts^16^. We validated these findings with another mouse strain, Swiss Webster which are known to be more resistant to *T. gondii* infection and as a consequence produce less tissue cysts (Supplementary Figure 2A). We found that ICI 56,780 reduced cyst burden by 86.1% at 1 mg/kg and 94.4% at 5 mg/kg compared to 80.1% for ATQ (Supplementary Figure 2B). All mice tested positive for the presence of ME49-RFP specific antibodies in their serum, confirming infection (Supplementary Figure 2C-E).

Altogether, these results demonstrate that ICI 56,780 is highly effective at reducing cyst burden in chronically infected mice, highlighting its potential as a standalone therapeutic candidate or as a lead scaffold for developing improved treatments for chronic toxoplasmosis.

## Discussion

In this study, we investigated the antiparasitic potential of two legacy 4(1*H*)-quinolones, ICI 56,780 and WR 243246, against *Toxoplasma gondii*. We focused on evaluating their effects on mitochondrial function and assessing their efficacy against both the acute and chronic stages of infection. Both compounds demonstrated potent inhibition of tachyzoite proliferation, including activity against ATQ-resistant strains, and they disrupted mitochondrial function by inhibiting electron transport and depolarizing the mitochondrial membrane potential. Notably, ICI 56,780 exhibited sub-nanomolar efficacy across multiple assays, including inhibition of mitochondrial activity and reduction of bradyzoite viability in both *in vitro* and *in vivo* derived parasites. In contrast, while WR 243246 was active against *in vitro* tachyzoites and bradyzoites, its effectiveness was markedly reduced against *in vivo* derived bradyzoites. This observation is consistent with the physicochemical profiling of ICI 56,780 and WR 243246, which showed that ICI 56,780 has better solubility and stability. These improved physicochemical properties likely contribute to ICI 56,780’s increased bioavailability compared to WR 243246. Impressively, *in vivo*, ICI 56,780 provided full protection in a lethal acute infection model at 10 mg/kg/day and significantly reduced brain cyst burden in chronically infected mice at low doses (1 and 5 mg/kg/day), outperforming the clinically approved ATQ. Together, these findings establish ICI 56,780 as a promising lead compound for treating both acute and chronic toxoplasmosis and highlight the importance of evaluating drug efficacy in physiologically relevant bradyzoite models. These results also support further medicinal chemistry optimization of both ICI 56,780 and WR 243246.

Mitochondrial function remains a critical vulnerability in *T. gondii* throughout both the tachyzoite and bradyzoite stages, making it an attractive target for therapeutic intervention in the chronic infection^14,17,18^. To assess whether these compounds target mitochondrial function via a distinct mechanism from ATQ, which inhibits the Q_o_ site of the mitochondrial *bc_1_* complex, we utilized ATQ^R^ *T. gondii* strains harboring mutations in the cytochrome *b* gene^7^. These strains were generated by McFadden et al and represent a valuable tool for characterizing the mitochondrial mechanism of action of ICI 56,780 and WR 243246. The emergence of *Plasmodium falciparum* resistance to ATQ has been well documented and is often linked to point mutations in the cytochrome *b* gene, which impairs drug binding and rapidly renders treatment ineffective^43^. The emergence of ATQ resistance is not considered a major clinical concern in *T. gondii*, largely because the parasite is not transmitted between humans, therefore, drug-resistant strains arising during treatment are unlikely to spread widely in the population^44^. However, between 2013 and 2017, six sulfadiazine-resistant strains were identified in Brazil, isolated either from patients with toxoplasmosis or from livestock intended for human consumption^45^. Therefore, analyzing resistant strains is important, not only for investigating the mechanism of action of mitochondrial inhibitors but also to monitor the potential emergence of clinically significant drug resistance. We attempted to generate ICI 56,780-resistant cell lines in three independent ENU mutagenesis experiments but were unsuccessful, as parasites failed to survive continued drug pressure after several passages.

Both ICI 56,780 and WR 243246 show comparable activity in ATQ^R^ and wild-type strains, suggesting that these quinolones inhibit mitochondrial function by engaging a site in the cytochrome *bc₁* complex distinct from the ATQ binding site. It has been shown that maintenance of mitochondrial membrane potential and ETC activity persists in bradyzoites and supports their long-term survival within tissue cysts^14^. Our findings that both ICI 56,780 and WR 243246 impair mitochondrial membrane potential and inhibit ETC activity further support this therapeutic approach. These results validate the mitochondrion as a druggable target during chronic infection and underscore the potential of mitochondrial inhibitors to overcome the limitations of current therapies, which fail to eliminate persistent tissue cysts. While our studies of mitochondrial inhibition focused on tachyzoites, additional work will be required to measure mitochondrial inhibition in bradyzoites, as it is difficult to obtain sufficient number of parasites to perform these analyses. It is also possible that quinolones affect other enzymatic functions or metabolic processes, as has been reported in viruses, bacteria, and cancer cells^20,46^.

Building upon the established efficacy of mitochondrial inhibitors against *T. gondii*, our study highlights the advantages of 4(1*H*)-quinolones, particularly ICI 56,780, relative to previously characterized ELQs such as ELQ-271 and ELQ-316. ELQs have shown potent *in vitro* activity and substantial reductions in brain cyst burden at doses of 5-25 mg/kg in murine models^17^. ICI 56,780 achieves comparable or greater efficacy at lower doses, reducing cyst burden by 77.7% at 1 mg/kg and up to 89.9% at 5 mg/kg. These findings underscore its enhanced potency and strong potential for improved therapeutic performance. This improved *in vivo* performance of ICI 56,780 may be due to its distinct chemical structure compared to ELQs, though it remains unclear whether and how these structural differences impact pharmacokinetic properties such as absorption, distribution, metabolism, and excretion, which determine bioavailability and tissue penetration. Moreover, potentially distinct binding interactions of 4(1*H*)-quinolones ICI 56,780 and WR 243246 within the cytochrome *bc₁* complex may confer advantages in overcoming resistance mechanisms associated with the Q_i_ site targeted by ELQs^17,47^. The ability of ICI 56,780 to maintain efficacy against ATQ-resistant strains further supports its potential as a robust chemotherapeutic agent.

## Conclusions

In summary, the 4(1*H*)-quinolone scaffolds ICI 56,780 and WR 243246 represent promising alternatives to existing mitochondrial inhibitors, combining potent anti-*T. gondii* activity with substantial medicinal chemistry optimization potential to improve physicochemical and pharmacokinetic properties. These findings support further exploration of both scaffolds for the treatment of acute and chronic toxoplasmosis.

ICI 56,780 exhibits exceptional potency across parasite stages, retains activity against ATQ-resistant strains, and achieves marked reductions in brain cyst burden at doses substantially lower than those required for ELQ-series compounds. Its broad activity profile, spanning inhibition of electron transport, disruption of mitochondrial membrane potential, tachyzoite killing, and direct activity against *in vivo*-derived bradyzoites, surpasses what is observed with current clinical agents such as pyrimethamine-sulfadiazine or ATQ, which do not eliminate encysted parasites. The ability of ICI 56,780 to maintain efficacy in the presence of *bc₁*mutations associated with ATQ resistance further highlights its potential as a next-generation mitochondrial inhibitor that may overcome limitations of present-day treatment regimens.

Together, these findings identify ICI 56,780 as a strong lead scaffold for further pharmacokinetic optimization and structure-guided medicinal chemistry refinement. Future work integrating mechanism-of-action studies and combination therapy approaches will be essential for defining the translational potential of this compound class. More broadly, this study reinforces the value of targeting parasite mitochondrial function, an essential activity maintained in bradyzoites, as a viable strategy for achieving more effective, possibly sterilizing, therapies for chronic toxoplasmosis.

## Materials and Methods

### Ethics Statement

All animal care and therapy studies were carried out in accordance with the NIH guidelines. The animal use protocol was reviewed and approved by the Institutional Animal Care and Use Committee (IACUC) of the University of Georgia. AUP# A2021 03-005-Y3-A4 and A2024 03-021Y2-A2.

### Cultures

*T. gondii* RH strain (all type I strains) was cultured using hTERT (human telomerase reverse transcriptase) cells with 1% bovine calf serum (BCS) and purified as described^48^. *T. gondii* ME49 (type II strain) were cultured using HFF (human foreskin fibroblasts) cells with 1% BCS. The ME49 parental strain was transfected with a Td-Tomato plasmid to create the ME49-RFP strain. Host cells were grown in Dulbecco’s modified Eagle medium supplemented with 10% BCS (hTERT) and 10% FBS (HFF). Cell cultures were maintained at 37°C with 5% CO_2_.

### Physiochemical property analysis

Physicochemical property experiments were performed by TCG Lifesciences (Kolkata, India). Determination of physiochemical properties and data analysis were performed as previously reported^49^.

### Synthesis of ICI 56,780 and WR 243246

#### General

Unless otherwise noted, all reagents and solvents were purchased from commercial sources and used without further purification. Tetrahydrofuran (THF) was distilled from benzophenone and sodium metal under an argon atmosphere immediately before use. Column chromatography was carried out using Sorbtec silica gel 60 Å (particle size 40-63 μm) and analytical thin layer chromatography was performed on 0.25 mm silica gel 60 F254 precoated plates from EMD Millipore. Microwave reactions were performed in an Anton Paar Monowave 400. The identity of all title compounds was verified via proton nuclear magnetic resonance (^1^H NMR) spectroscopy and liquid chromatography with low-resolution mass spectrometry detection (LRMS). ^1^H NMR spectra were recorded at ambient temperature on a Bruker 500 or 700 MHz spectrometer. All ^1^H NMR experiments are reported in δ units, parts per million (ppm) downfield of trimethyl silane (TMS) and were measured relative to the residual proton signals of chloroform (δ 7.26) and dimethylsulfoxide (δ 2.50). Data for ^1^H NMR are reported as follows: chemical shift (δ ppm), multiplicity (bs = broad singlet, s = singlet, d = doublet, dd = doublet of doublets, dt = doublet of triplets, ddt = doublet of doublet of triplets, t = triplet, tt = triplet of triplets, q = quartet, hept = heptet, m = multiplet), coupling constant (Hz), and integration. NMR data was analyzed by using MestReNova Software version 12.0.3-21384. Low resolution mass spectra (LRMS) were acquired on an Agilent 1260 Infinity HPLC with a thermostatted well-plate autosampler and diode array detector hyphenated to an Agilent G6125B single quadrupole mass spectrometer with electrospray ionization. ionization. At a flow rate of 0.6 mL/min, samples were analyzed on an Agilent ZORBAX RRHT StableBond-C18 column (1.8 µm, 2.1 x 50 mm, part no: 827700-902) with the following HPLC method: 0 min 90/10 A/B, 3.5 min 10/90 A/B, 5.5 min 10/90 A/B, 6.0 min 90/10 A/B wherein A is water (+0.1% formic acid) and B is acetonitrile (+0.1% formic acid). All final compounds are >95% pure by HPLC analysis.

#### Synthesis of ICI 56,780

ICI 56,780 was synthesized as previously reported^28^. The crude material was purified via recrystallization in dichloromethane to afford a white solid. ^1^H NMR (500 MHz, DMSO-d6) δ 12.11 (s, 1H), 8.49 (s, 1H), 7.88 (s, 1H), 7.31 (dd, J = 7.8 Hz, 2H), 7.06 (s, 1H), 7.02 – 6.94 (m, 3H), 4.41 (s, 4H), 3.73 (s, 3H), 2.62 (t, J = 7.6 Hz, 2H), 1.53 (p, J = 7.4 Hz, 2H), 1.29 – 1.22 (m, 2H), 0.84 (t, J = 7.3 Hz, 3H). LRMS-ESI (*m/z*): [M + H]^+^ 396.2; retention time: 3.92 min. TLC (5% methanol in dichloromethane) Rf 0.41 (UV, 254 nm, 280 nm).

#### Synthesis WR 243246

WR 243246 was synthesized as previously reported^27^. The crude material was purified via recrystallization in dichloromethane to afford a yellow solid. ^1^H NMR (500 MHz, Chloroform-d) δ 9.77 (s, 1H), 8.49 (d, J = 2.1 Hz, 1H), 8.14 – 8.04 (m, 2H), 7.50 (d, J = 2.1 Hz, 1H), 7.35 (dd, J = 8.4, 2.1 Hz, 1H), 7.25 (s, 1H), 4.20 – 4.09 (m, 1H), 3.79 – 3.63 (m, 2H), 3.27 – 3.19 (m, 1H), 3.17 – 3.07 (m, 1H). LRMS-ESI (*m/z*): [M + H]^+^ 391.6; retention time 3.27 min. TLC (5% methanol in dichloromethane) Rf 0.38 (UV, 254 nm, 280 nm).

### *In vitro* dose response curves

Experiments with *T. gondii* tachyzoites were carried out with parasites expressing a td-Tomato red fluorescent protein (RFP)^50,51^. Parasites were purified by passing them through a 25-gauge needle, followed by filtration through a 5 μm filter. Human fibroblasts were cultured in 96 well black plates for 48 (hTERT) or 120 (HFF) hours prior to the addition of 4,000 fluorescent tachyzoites/well. Compounds were tested at five different concentrations to generate dose-response curves for ICI 56,780 and WR 243246, while ATQ was tested at ten different concentrations. Two technical replicates per concentration were used, and the experiment was repeated in three independent biological replicates. Fluorescence values were measured for up to 7 days and both excitation (544 nm) and emission (590 nm) were read from the bottom of the plates in a BioTek Synergy H1 Hybrid plate reader^16^. For the dose-response curves of the ATQ^R^ strains which are not fluorescent, parasites were allowed to grow for 7 days before staining the plates with crystal violet and reading the absorbance of the wells at 590 nm using a BioTek Synergy H1 plate reader. The EC_50_s were calculated using Graph Prism software version 10.

### Cytochrome c reduction assay

The assay was carried out using a previously published protocol^52^. Five µg of protein from a *T. gondii* mitochondrial enriched fraction (P2) was used for each technical replicate (3 in total). After addition of reaction buffer (0.3 mM succinate, 0.3 mM KCN, 10 mM HEPES, pH 7.5), with succinate as the substrate and each compound at 100 nM, the reaction was initiated by addition of cytochrome c (0.1 mM final concentration). The absorbance of reduced cytochrome c was measured at 550 nm using a BioTek Synergy H1 plate reader for 10 min with shaking at 30°C. The activity of each fraction was then normalized to protein concentration as determined by the BCA assay.

### Mitochondrial membrane potential

The mitochondrial membrane potential was measured by the safranine O method according to published protocols^36,37^. Freshly lysed parasites were collected and filtered through an 8 µm filter to remove host cell debris. The parasites were washed twice with BAG (116 mM NaCl, 5.4 mM KCl, 0.8 mM MgSO_4_ 7H_2_O, 50 mM HEPES, 5.5 mM Glucose, pH 7.3) and resuspended at 1 x 10^9^parasites per ml. An aliquot of 50 µl of the parasite suspension (5 x 10^7^ cells) was added to a cuvette containing Safranine O (2.5 µM) and succinate (1 mM) in 2 ml of reaction buffer (125 mM sucrose, 65 mM KCl, 10 mM HEPES-KOH pH 7.2, 1.0 mM MgCl_2_, and 2.5 mM K_3_PO_4_ pH 7.2).

The cuvette was placed in a Hitachi F-7000 fluorescence spectrophotometer and digitonin (10 µM) was added to selectively permeabilize the plasma membrane. After equilibration, ADP (10 µM final) was added to stimulate oxidative phosphorylation followed by the compounds at 2.5 µM (ATQ) and 1 µM (ICI 56,780 and WR 243246). The change in mitochondrial membrane potential was quantified as the difference between the maximum fluorescence (after compound addition) and the minimum fluorescence (following ADP stabilization and prior to compound addition). Statistical analysis was performed using two-way ANOVA in GraphPad Prism 10.

### Acute *in vivo* infection in mice

Experiments were carried out as described previously^53^ using 100 *T. gondii* tachyzoites of the RH td-Tomato strain (RH-RFP) to infect Swiss Webster mice. Drugs were dissolved in 10% Kolliphor HS-15 and were inoculated intraperitoneally (i.p.) starting 6 hours after infection and administered for 10 days. The survival of mice was monitored along with their weight. Mice were sacrificed if they lost over 20% of their body weight. Surviving mice were challenged on day 30 post-infection with 10,000 RH-RFP tachyzoites. Blood was collected at 28 dpi (before challenge) and at the time of euthanasia (3 weeks post challenge) to analyze the presence of *T. gondii* antibodies. To test for antibodies, we performed western blots with RH-RFP lysates and probed the membrane with serum collected from the surviving mice. Protein extraction and western blot conditions were done as previously^15^. After blocking with 5% milk in PBS-T (0.1% tween 20 in PBS), the membranes were washed 3 times for 5 min with PBS-T and attached to the BioRad multiscreen tool. The RH-RFP lysate was run across the entire gel to allow testing of different sera (antibodies) on the same membrane. We incubated the membrane with serum (used as the primary antibody) for 1 h in the multiscreen tool and then washed the membrane five times for 3 min each while still in the multiscreen, followed by one additional wash after removal. Then we used IRDye 800CW goat anti-mouse IgG secondary antibody and incubated for 1 h before washing three times with PBS-T and imaging the blot on a LI-COR Odyssey CLx at 800 nm.

### *In vitro* bradyzoite differentiation and immunofluorescence assay

Type II ME49 was differentiated using compound 1 (4-[2-(4-fluorophenyl)-5-(1-methylpiperidine-4-yl)-1H-pyrrol-3-yl]pyridine) as previously described^54^. Confluent HFF monolayers on glass coverslips (preincubated for 2 hours with 3 µM compound 1, as described^39^) were infected with 5 x 10^3^ tachyzoites of the ME49 strain. The cells were allowed to grow and differentiate in a 37°C incubator with ambient CO_2_ for 3 days. The cells were then incubated with drugs at 3 times their EC_50_ for an additional 4 days at ambient CO_2_. After 7 total days of incubation the cells were fixed with 3% paraformaldehyde for 10 min, permeabilized with 0.25% triton X-100 for 10 min, and then blocked in 3% bovine serum albumin (BSA) for 1 hour. The coverslips were then incubated in Dolichos biflorus agglutinin (DBA) rhodamine (1:2,000)^55^, anti-BAG1 rabbit (1:1,000), and anti-SAG1 rat (1:1,000) antibody (both a kind gift from Vern Carruthers) for 2 hours. Then the secondary anti-rat 488 was used to visualize the SAG1 antibody. The cells were then mounted onto microscope slides with fluoromount and DAPI. The slides were placed in the fridge for a minimum of 24 h before imaging on a Delta Vision microscope. The vacuole sizes were measured using ImageJ software.

### *In vitro* bradyzoite viability assay

The protocol for *in vitro* viability assay was adapted from Zwicker et al.^56^. Confluent HFF monolayers (preincubated for 2 h with 3 µM compound 1) were infected with 5 x 10^3^ tachyzoites of the ME49 (type II) strain. After 24 h, compound 1 was removed to avoid toxicity to the host cells. The cells were allowed to grow and differentiate in a 37°C incubator with ambient CO_2_ for 3 days. The cysts were then incubated with drugs at 30 nM for an additional 4 days at ambient CO_2_. After 7 days of differentiation the cysts were collected by scraping the monolayers, syringe lysing the parasites, and treatment with acid/pepsin (170 mM sodium chloride, 60 mM hydrochloric acid, and 0.1 mg/ml pepsin) for 30 min in a 37°C water bath. The cyst suspension was neutralized with 94 mM NA_2_CO_3_ and enumerated. 10,000 bradyzoites were plated per well in a 6-well plate and allowed to grow undisturbed in a 37°C incubator with 5% CO_2_ for 16 days. At this time, cultures were fixed with 100% ethanol and stained with 2.5X crystal violet. Plaques were quantified using a light microscope using the 4X objective and were enumerated and normalized as a percentage of the plaques formed by the DMSO control.

### Isolation and *ex vivo* assay of *in vivo-*derived bradyzoites

Brains of mice chronically infected with ME49 tachyzoites i.p. were collected 28 dpi, homogenized with 7 strokes in a glass homogenizer, and then enumerated on a Delta Vision microscope (protocol adapted from^42^). For 12 well plate assays, 25 cysts per plate were placed in a tube for acid/pepsin (170 mM sodium chloride, 60 mM hydrochloric acid, and 0.1 mg/mL pepsin) treatment for 10 min in a 37°C water bath. The cyst suspension was then neutralized with 94 mM sodium carbonate (NA_2_CO_3_). The liberated bradyzoites were then equally distributed into a 12 well plate. For the growth assay the cells were grown in the presence of compounds or DMSO for 12 and 14 days undisturbed. For the viability assays the cells were treated with compounds or DMSO for 4 h before being washed with 1X PBS and then grown for 14 days in drug-free media. After 14 days the cells were fixed with 100% ethanol and stained with 2.5 X crystal violet. We also adapted this assay to a 96-well plate format to generate dose-response curves. In this format, 500 cysts were seeded per plate, and both growth and viability assays were performed as described above. Plaques from the 12-well plates were quantified on a light microscope using the 4X objective and the size of the plaques was measured using Fiji (ImageJ) software. For the 96-well plate assays we measured the absorbance at 590 using a BioTek Synergy H1 plate reader.

### Chronic *Toxoplasma* infection in mice

CBA/j mice (Jackson Laboratory) were infected i.p. with a 200 μL suspension of 1,000 tachyzoites of the ME49-RFP strain *T. gondii* (type II genotype). About 15% of the mice succumbed to the acute infection before 28 days post infection. At 4 weeks post infection the brain of 2 mice were collected to check for the presence of cysts and to confirm that the mice entered the chronic stage. At this time the remaining mice were injected i.p. daily for 16 consecutive days with 200 μL solvent (10% kolliphor HS-15) or compounds dissolved in 10% HS-15. ATQ was administered at 5 mg/kg and ICI 56,780 was administered at 1 mg/kg or 5 mg/kg. Mice were euthanized humanely 2 weeks after the final treatment. The mouse brains were collected in 1 mL sterile PBS, minced with scissors and homogenized using a glass homogenizer. Four 20 μL samples (80 μL total or 8% of sample) of each brain homogenate was placed on a glass microscope slide. Cysts were enumerated in a Delta vision fluorescence microscope.

### Statistical Analysis

Experimental data are expressed as the mean values ± standard error of the mean (SEM) from at least 3 independent biological replicates unless indicated otherwise. Statistical analyses were performed using two-way Anova statistical analysis using GraphPad PRISM version 10. A P-value of < 0.05 was considered statistically significant.

## Acknowledgements

We thank Vern Carruthers for the BAG1 and SAG1 antibodies, and John Boothroyd for providing the ATQ-resistant strains. We thank Zhu-Hong Li for generating the ME49-RFP cell line. This work was supported by NIH grants R01AI169846 (SNJM) and R01AI144464 (RM). MAS was partially supported by the T32AI060546 predoctoral training grant.

