## Supplemental Figures and Tables for "Legacy 4(1*H*)-quinolone scaffolds activity against acute and chronic *Toxoplasma gondii* infection"

**Supplementary Tables and Figures**

**Supplementary Table 1.** Cytotoxicity in HFF fibroblasts from AlamarBlue assay. Three independent biological replicates with two technical replicates per concentration.

| Concentration | % Inhibition (± standard deviation) | | |
|---|---|---|---|
| | ATQ | ICI 56,780 | WR 243246 |
| 0.1 nM | 4.72 ± 1.14 | 5.86 ± 3.94 | 3.60 ± 3.66 |
| 0.5 nM | 6.81 ± 0.96 | 5.49 ± 4.70 | 7.05 ± 4.42 |
| 1 nM | 5.92 ± 2.25 | 4.73 ± 3.46 | 5.61 ± 4.79 |
| 100 nM | 6.37 ± 2.29 | 7.18 ± 3.76 | 6.17 ± 3.32 |
| 500 nM | 7.85 ± 3.74 | 7.31 ± 4.36 | 6.34 ± 5.27 |
| 1 µM | 7.58 ± 2.71 | 8.09 ± 2.65 | 7.63 ± 2.23 |
| 5 µM | 7.37 ± 1.55 | 6.04 ± 1.58 | 6.89 ± 1.91 |
| 10 µM | 8.90 ± 1.49 | 5.97 ± 0.86 | 6.63 ± 1.94 |
| 25 µM | 9.62 ± 3.35 | 5.17 ± 2.23 | 6.89 ± 1.80 |
| 50 µM | 8.32 ± 3.72 | 6.91 ± 0.25 | 3.76 ± 2.07 |

**Supplementary Table 2.** Cytotoxicity in BV-2 Microglial cells from AlamarBlue assay. Three independent biological replicates with two technical replicates per concentration.

| **Concentration** | **% Inhibition (± standard deviation)** | | |
|---|---|---|---|
| | **ATQ ($CC_{50}$=33.24 µM)** | **ICI 56,780 ($CC_{50}$>50 µM)** | **WR 243246 ($CC_{50}$=82.6 µM)** |
| 0.1 nM | 8.97 ± 0.94 | 4.12 ± 2.91 | 3.01 ± 1.57 |
| 0.5 nM | 10.1 ± 5.36 | 8.73 ± 7.63 | 5.03 ± 4.21 |
| 1 nM | 6.55 ± 1.04 | 6.16 ± 3.44 | 2.76 ± 5.99 |
| 100 nM | 9.38 ± 6.24 | 8.73 ± 4.91 | 3.47 ± 4.72 |
| 500 nM | 7.97 ± 5.92 | 10.6 ± 4.57 | 9.42 ± 3.00 |
| 1 µM | 6.98 ± 3.88 | 9.98 ± 4.52 | 9.16 ± 4.31 |
| 5 µM | 34.1 ± 13.6 | 9.36 ± 7.76 | 8.22 ± 1.31 |
| 10 µM | 34.4 ± 15.6 | 11.7 ± 9.71 | 7.33 ± 6.88 |
| 25 µM | 43.5 ± 0.79 | 15.3 ± 1.70 | 20.8 ± 1.11 |
| 50 µM | 55.1 ± 9.60 | 27.1 ± 14.7 | 40.2 ± 11.6 |

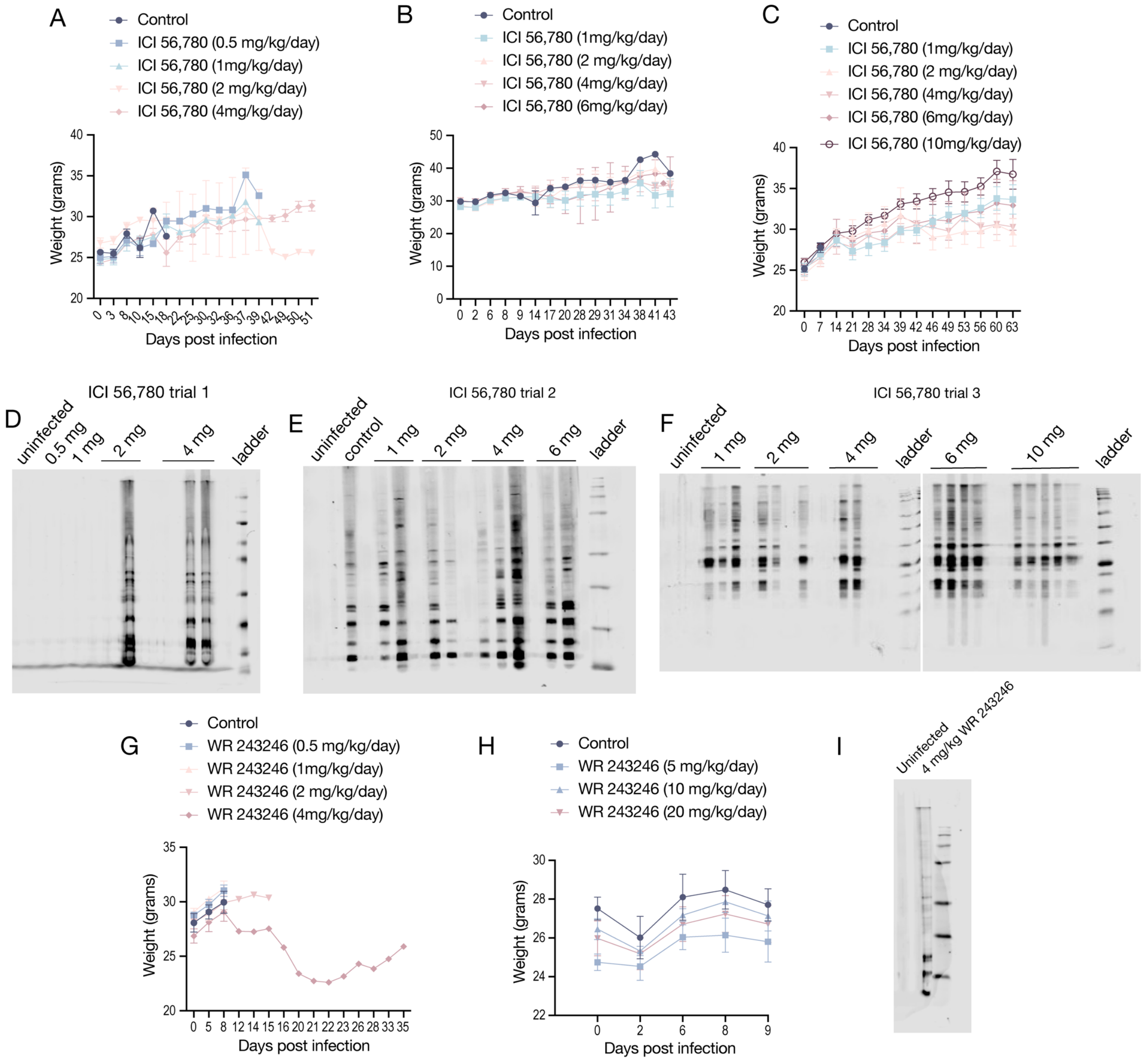


**Supplementary Figure 1.** Additional data for the *in vivo* acute infections in mice. A-C. Weight curves for trials 1-3 of the *in vivo* acute infection of mice treated with ICI 56,780. D-F. Serum tests for RH-RFP total protein lysate for serums from trials 1-3 from treated mice with ICI 56,780 before challenge (28 dpi). G-H. Weight curves for trial 1 and 2 of the *in vivo* acute infection for mice treated with WR 243246. I. Pre-challenge serum test (28 dpi) for the one surviving mouse during trial 1 treated with 4 mg/kg dose of WR 243246.

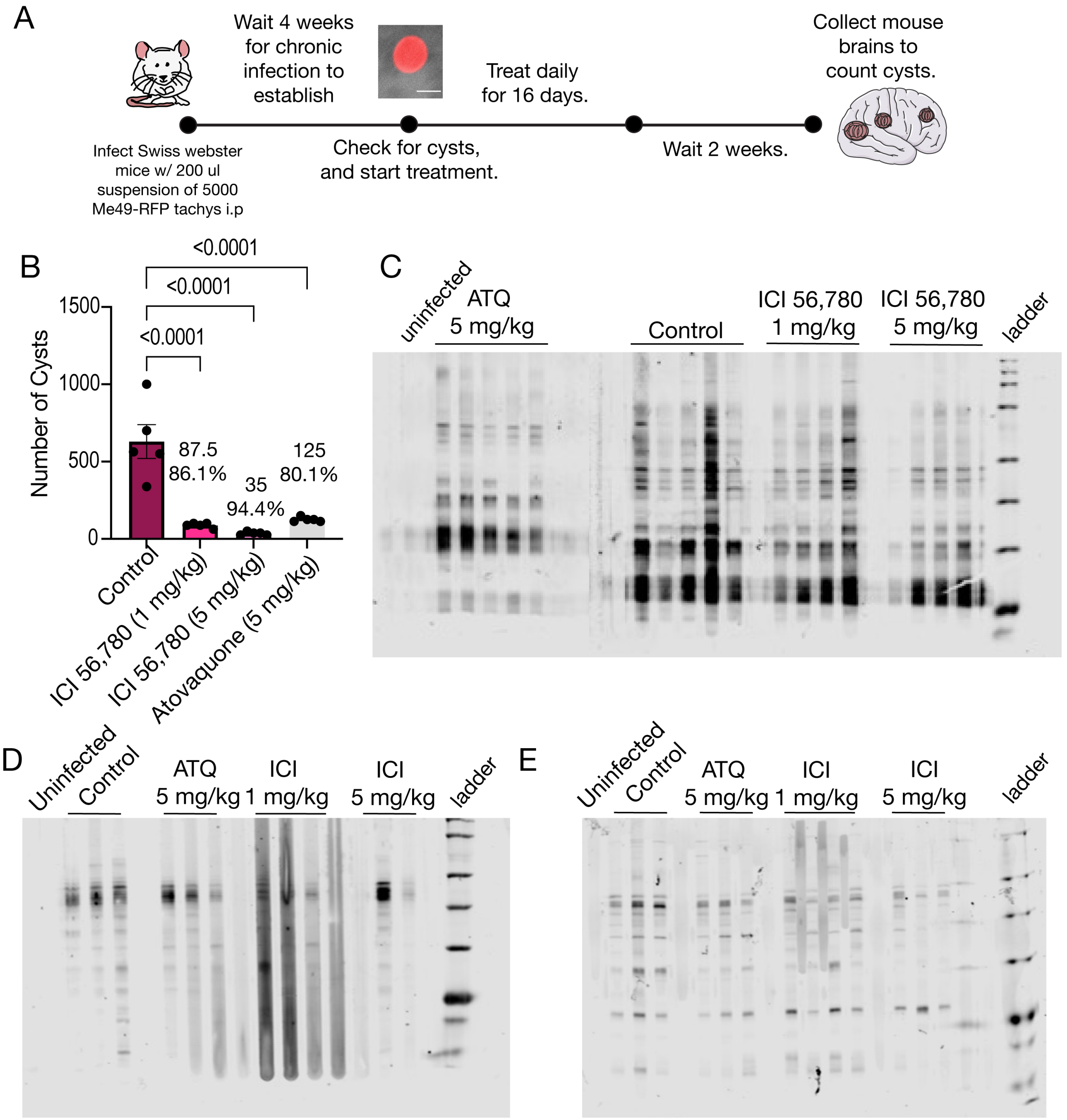


**Supplementary Figure 2.** Treatment with ICI 56,780 of chronically infected Swiss Webster mice. Scheme of the protocol followed for *in vivo* chronic infection with female Swiss Webster mice. Mice were infected with 5,000 Me49-RFP tachyzoites i.p. Four weeks post-infection 2 mice were euthanized, and their brains collected to check for the presence of cysts. Daily treatment i.p was initiated for 16 days with ICI 56,780 and ATQ at the indicated doses. B. Quantification of the number of cysts. Each dot represents a mouse (5 mice per group). C. Serum analysis from Swiss webster mice chronically infected with Me49-RFP parasites. D. Serum analysis using Me49-RFP total protein lysate from trial 1 of the chronic infection with CBA/j mice. E. Serum analysis using Me49-RFP total protein lysate from trial 2 of the chronic infection with CBA/j mice.
